# Aminooxadiazolyl kainic acid reveals that kainic acid receptors contribute to astrocytoma glutamate signaling

**DOI:** 10.1101/2021.01.16.426948

**Authors:** Mitra Sadat Tabatabaee, Zhenlin Tian, Julien Gibon, Frederic Menard

## Abstract

The excitatory neurotransmitter glutamate triggers a Ca^2+^ rise and the extension of processes in astrocytes. Our results suggest that kainic acid receptors (KAR) can independently initiate glutamate signaling in astrocytoma U118-MG cells. The natural product kainic acid triggered glioexcitablity in cells and was inhibited by the KAR antagonist CNQX, but its activity was lower than glutamate on KARs. We created a new heteroaryl kainoid based on rational design: aminooxadiazolyl kainic acid **1** (AODKA). AODKA induced a larger calcium influx and a faster processes extension than kainic acid in U118-MG cells. AODKA is a new tool to study KAR activity in the nervous system.

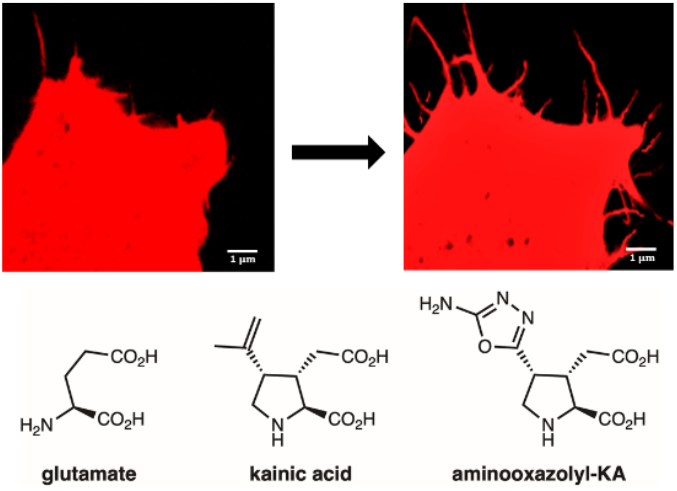

---

Ionotropic glutamate receptors (iGluRs) are ligand-gated ion channels activated by glutamate at excitatory synapses; they are necessary for signal transduction in the central nervous system (CNS).^1^ The glutamate released by a pre-synaptic neuron has been shown to also activate glutamate receptors in astrocytes.^2-4^

In the brain, a synapse is typically ensheated by the process of an astrocyte.^5^ It is now established that astrocytes processes extension and retraction at synapses play an essential role in neuronal development, synaptic plasticity, energy support, neuro-glial vascular integration, and healthy cognitive function.^6,7^ In the astrocytic network, glutamate initiates a form of cellular excitation via an intracellular calcium rise ([Ca^2+^]i) leading to long-range Ca^2+^ waves, i.e., glioexcitability.^2,3,8,9^ This local [Ca^2+^]i rise leads to actin remodeling and is believed to be the mechanism driving the motility of astrocyte’s processes at synapses (Fig. 1).^3,5,10^ Thus, it is important to identify which glutamate receptors govern glioexcitability to understand this phenomenon in health and disease.

**Figure 1.**
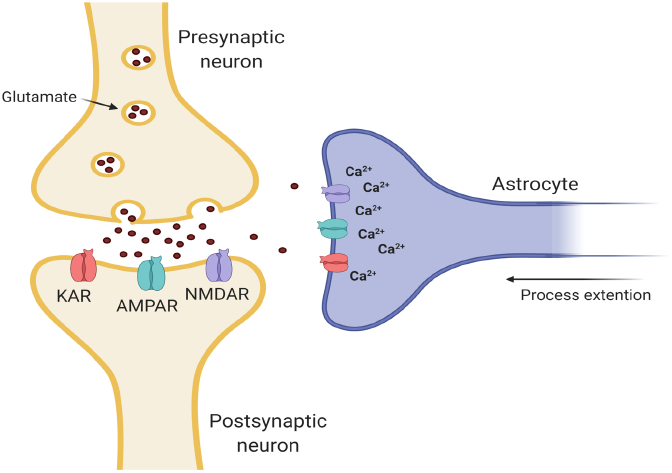
Neurons and astrocytes express ionotropic glutamate receptors that are also activated by NMDA, AMPA, or kainic acid (KA). Upon binding of glutamate or selective agonists, the receptors allow an influx of calcium ions that initiates intracellular events such as depolarizations in postsynaptic neurons, and process extension in astrocytes.

In mammals, iGluRs are categorized based on their preferential activation by three agonists: α-amino-3-hydroxy-5-methyl-4-isoxazolylpropionic acid (AMPA), *N*-methyl-D-aspartate (NMDA), and kainic acid (KA).^11^ Alternatively, metabotropic glutamate receptors (mGluRs) are another family of proteins that can release calcium from intracellular stores, and glutamate is their only known activating ligand.^12,13^

Among iGluRs, KARs have been the least explored and their relevance to pathophysiology is largely unknown. This is partially due to the low abundance of KARs compared to other channels, combined to the lack of specific agonists and antagonists for this receptor subtype. Consequently, KARs’ function in neuronal network may have been underestimated.^14^

In cultured astrocytes, AMPA and KA have been shown to initiate glioexcitability to the same level as glutamate (NMDA is ineffective).^3,10,15^ While the role of AMPA receptors and mGluRs in astrocytic glutamate signaling has been examined, the contribution of kainate receptors (KARs) is still unexplored.^3,10,13,16^ In our work with astrocyte model cells U118-MG, we found serendipitously that KARs were the only glutamate receptors responsible for glioexcitability, and that KA is not as effective as glutamate to initiate the response. To show unequivocally the independent role of KARs in glioexcitability, we designed a novel KAR agonist: aminooxadiazolyl kainic acid (AODKA, **1**), based on the hypothesis that a heteroaryl analog with enhanced π–π stacking ability would be more potent than KA.^17^ Below we show that: KARs are sufficient to initiate glioexcitability, KA is not as effective as glutamate, and AODKA is highly effective at activating KARs.

## Kainic acid receptors can initiate filopodiagenesis independently

The exact mechanism of the [Ca^2+^]i rise that leads to filopodia extension (glioexcitability) in astrocytes upon exposure to glutamate is largely unknown.^3,15^ Different subtypes of mGluRs or iGluRs have been reported to initiate this cellular response.^3^

We selected the astrocytoma cell line U118-MG as a model to study glutamate receptors participate in its glutamate signaling. Well established GluRs agonists that initiate glioexcitability (e.g., glutamate, KA and AMPA) were examined if they can initiate response in U118-MG cells. Cells responded morphologically to glutamate and KA, by forming new processes, and extending the existing ones farther (Fig. 2.A). In contrast, AMPA did not cause such a response, which implies that filopodiagenesis is due to the activity of KARs and/or mGluRs, but not AMPARs. To confirm that the response we observed in exposure to KA is an is not affected by cellular viability, variation between batches or treatment conditions, we monitored the same cells while beeing treated with AMPA and then being exposed to KA (Fig. 2B and 2C). Given that KA is selective for KARs and does not activate mGluRs, KARs are appeared to be solely responsible for the phenotype in U118-MG cells. We tested the effect of another kainoid known to bind selectively to KARs: phenylkainic acid (Ph-KA), to confirm further our observation.^18,19^ Ph-KA caused the same response as KA (Fig. 4). To verify that KARs are directly involved, we treated the cells with CNQX-KARs’ most selective antagonist,.^18^ CNQX shut down all processes extension not only in exposure to KA and Ph-KA but also to glutamate (Fig. 4B). The above observations suggest that KARs play a key role in initiating glioexcitability of glutamate in U118-MG cells.

**Figure 2.**
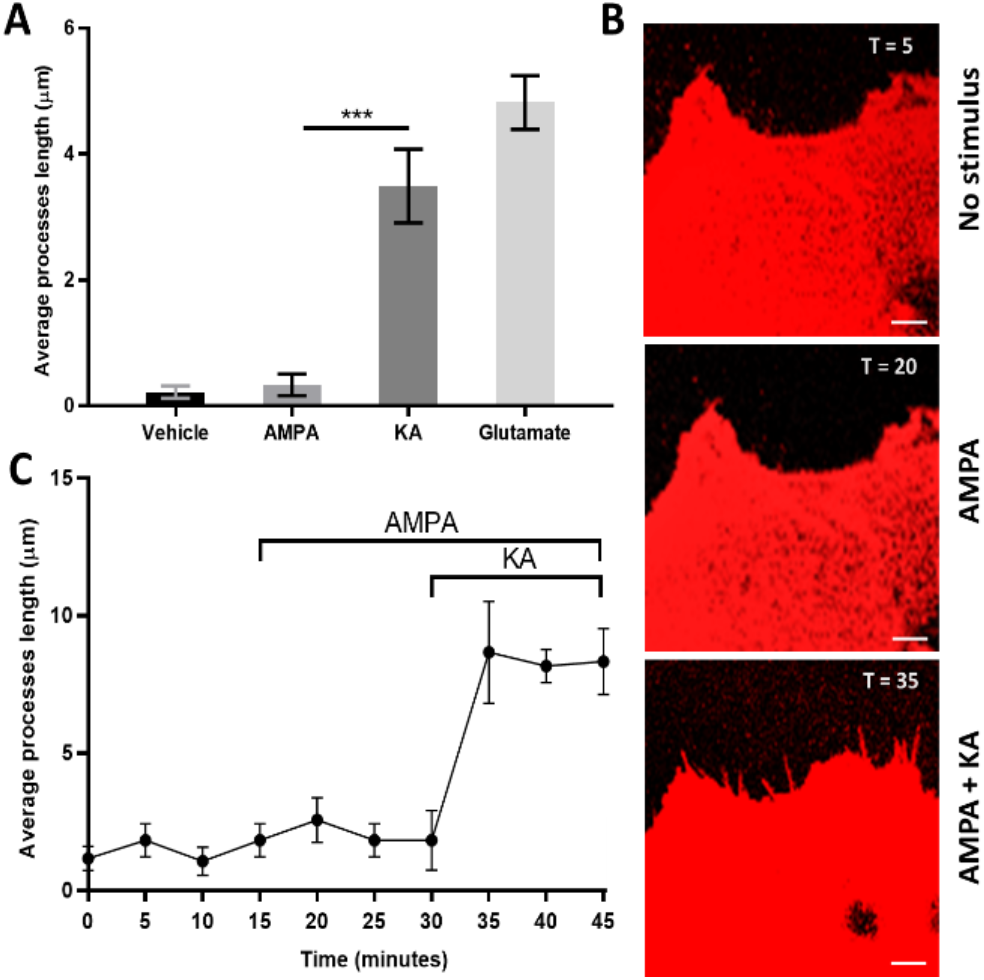
Kanic acid, but not AMPA, triggers filopodiagenesis in astrocytoma U118-MG cells. (**A**) Cells extend their processes significantly in response to 100 μM of glutamate and KA. AMPA does not have the same effect. Change in length (Δ) was calculated by subtracting the length of a filopodium immediately prior to stimulation from that of the same filopodium after 20 minutes. Results are presented as mean ± SEM. The change in length is the average data of 5 cells, each from a separate culture dish; measurements of at least 20 filopodia were averaged for each cell (n = 5). Comparisons was done with one-way ANOVA folllowed by a post hoc Tukey’s multiple comparison ***p = 0.0001. (**B**) Representative confocal micrographs of a cell transfected with mCherry-LifeAct7, before and after exposure to 100 μM of AMPA and then to KA. Scale bar is 1 μm. (**C**) Averaged length of processes during a time-lapse imaging response to stimulation of 100 μM AMPA added after 15 minutes and KA added after 30 minutes to the same culture dish; n = 3, mean ± SEM.

Since glioexcitability is associated with calcium influx in astrocytes, calcium imaging experiments were performed with the intracellular calcium fluorescent dye Fluo4-AM. Stimulation of the cells with 100 μM solutions of KA and Ph-KA showed weaker [Ca^2+^]i spikes than with glutamate (2.3- and 3.6-fold, respectively, Fig. 4C). In line with this result, KA and Ph-KA also induced a delayed response compared to glutamate when the filopodiagenesis was recorded by time-lapse imaging (Fig. 4D).^3^ These unanticipated differences suggest that KARs in U118-MG cells are more phenotypically sensitive to glutamate stimulation than to KA and Ph-KA (Fig. 4C). This lower phenotypic activity of KA may have led to the contribution of KARs being underestimated in the CNS (kainic acid is the only broadly used commercial kainoid). To unequivocally demonstrate the independent contribution of KARs in glutamate signaling of U118-MG cells, we needed a more effective molecular tool than KA and PhKA. Thus, we designed AODKA as a more potent agonist for KARs.

## Aminooxadiazolyl kainic acid

Kainoids are a group of nonproteinogenic amino acids that were initially isolated from the marine algae *Digenea simplex*.^17^ The parent natural product, kainic acid, has been widely used as a chemical tool in neurobiology.^17,20-22^ Below is described the synthesis of AODKA as lead compound for a novel class of heteroaromatic KA analogs (Scheme 1).

**Scheme 1.**
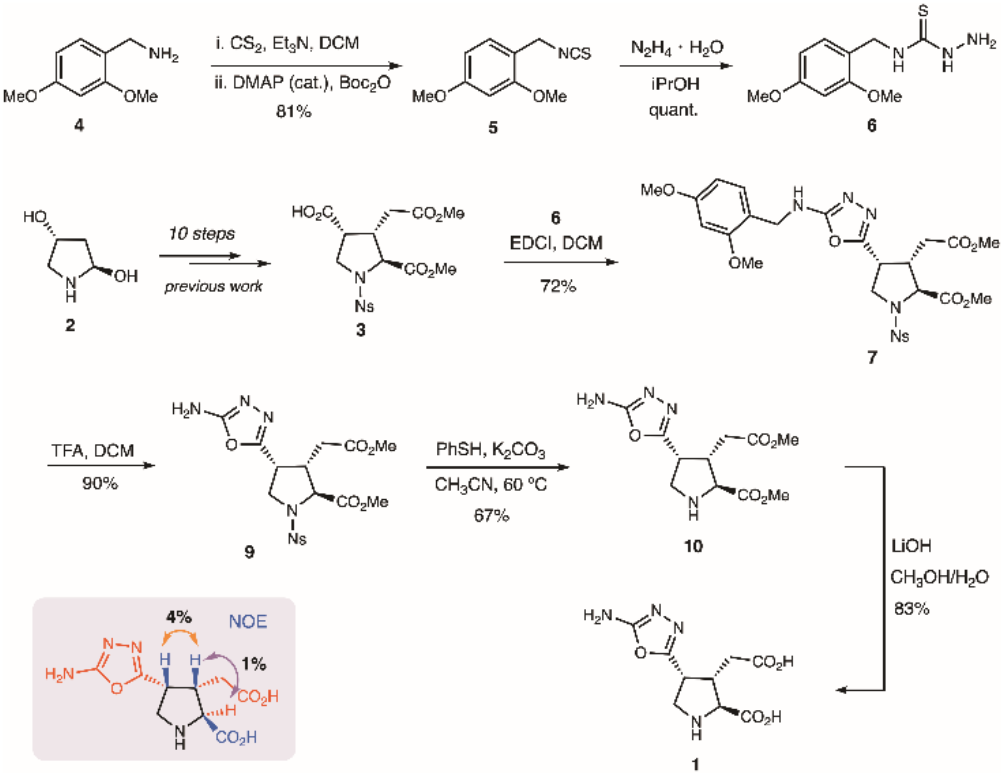
Synthesis of aminooxadiazolyl kainic acid (**1**).

Based on our practical synthesis of kainoid analogs, we designed a C4-heteroaryl kainoid that was expected to be more potent than its parent natural product kainic acid.^17,19,20^ Our previous SAR analysis^12^ suggested that C4-aryl kainoids are highly potent KAR agonists, mostly due to the C4 substituent participating in π-π stacking with a key phenol side chain within the binding pocket (Tyr448, Figure 3). Aminooxazolyl kainoid **1** was selected for two main reasons: (1) its electron-deficient quadrupole would complement and enhance π-π stacking interactions with Tyr488, and (2) the 2-amino substituent would allow easy attachment of different molecular cargo to create new chemical biology tools (e.g., fluorescent tags, biotin, photoswitch, etc.).

**Figure 3.**
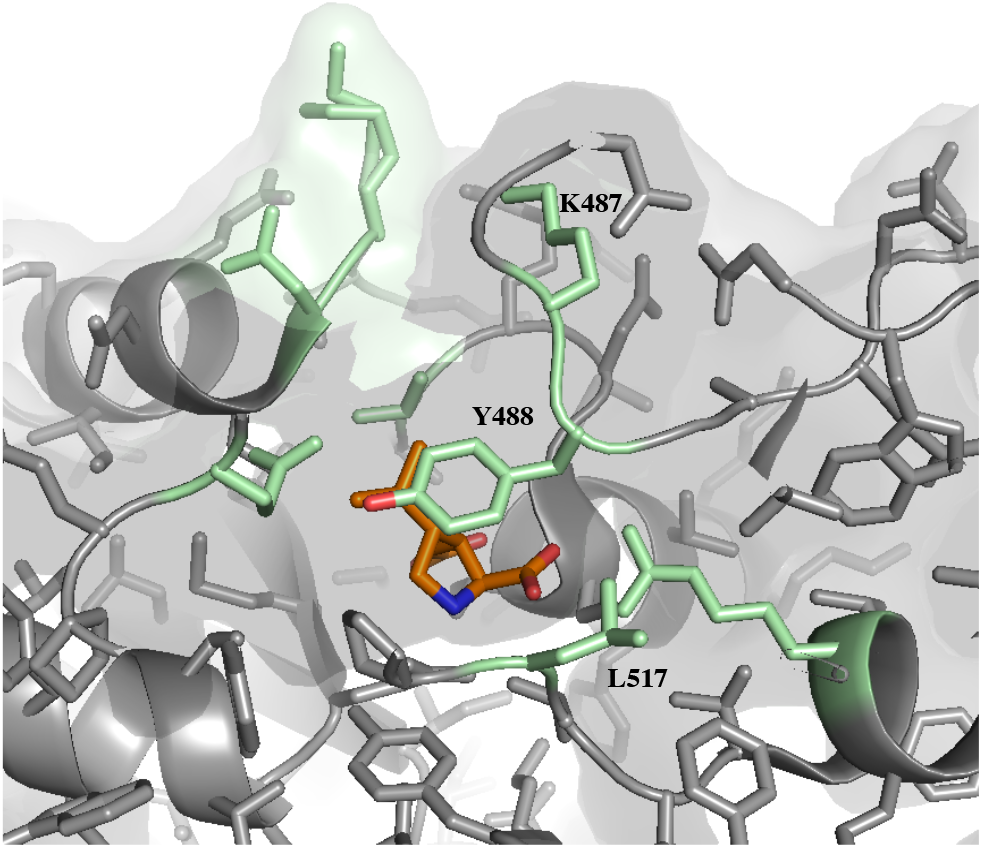
Kainic acid’s alignment can make a π-π interaction with Tyr488 in GluK receptors. Top views of kainic acid bound to GluK1 (PDB: 3c33). This similar binding mode has been observed on GluK1-5.

**Figure 4.**
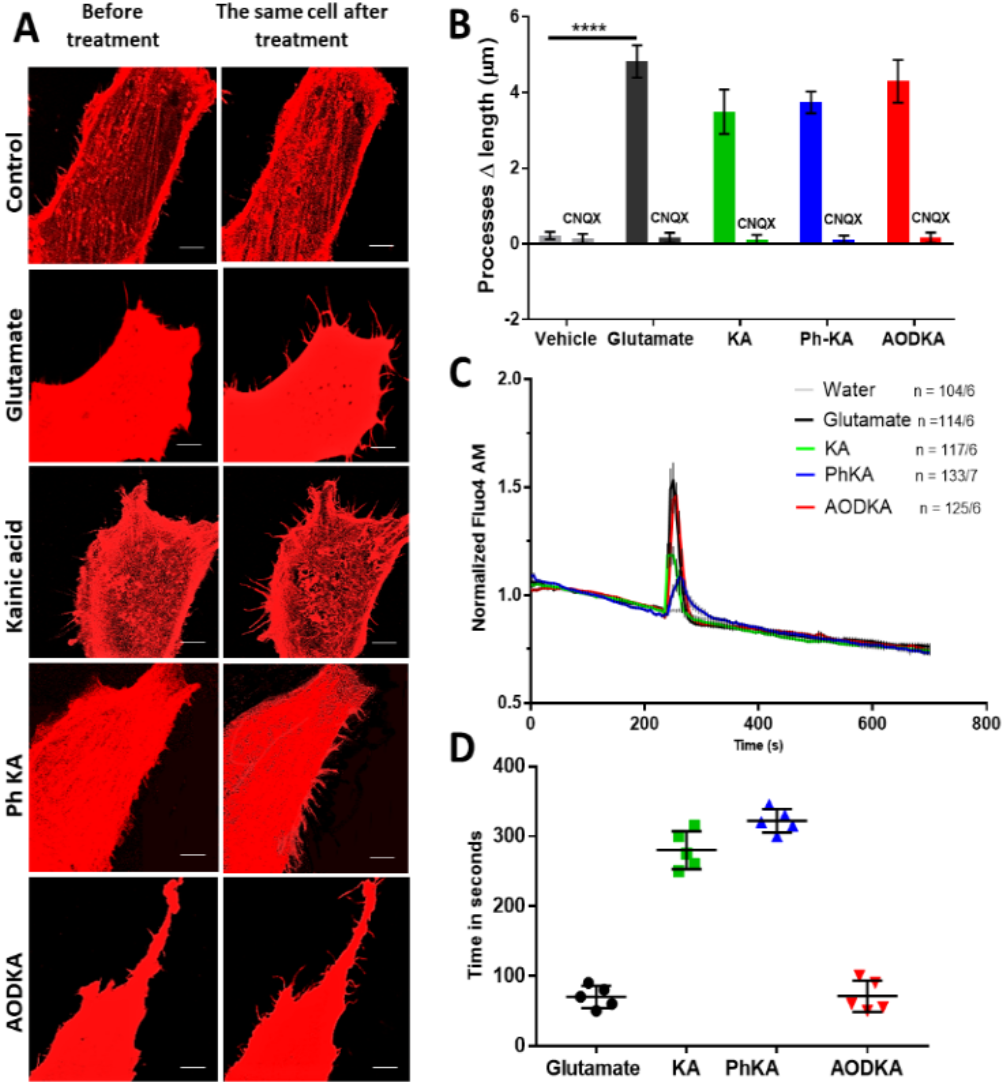
AODKA causes filopodiagenesis and an intracellular calcium rise in U118-MG astrocytoma cells similar to glutamate. (A) Representative confocal micrographs of cells transfected with mCherry-LifeAct7, before and after exposure to 100 μM Glu, KA, Ph-KA, or AODKA. Scale bar is 5 μm. (B) Change in a cell’s processes length when stimulated with 100 μM glutamate and kainoids compared to cells pre-treated with 10 μM CNQX antagonist for 20 minutes. Sidak’s multiple comparison post hoc (two-way ANOVA); n = 5; ****p < 0.0001. (C) Normalized fluorescence of Ca^2+^ imaging dye Fluo4-AM in U118-MG cells stimulated with 100 μM kainoids or glutamate (n = 104/6). (D) Time elapsed before the initiation of process extension after chemical stimuli; AODKA is 3-4 times faster than analogs KA or PhKA. n=5. mean ± SEM.

The synthesis of novel aminooxadiazolyl kainoid **1** began with the preparation of *N*-protected thiosemicarbazide **4** from dimethoxybenzylamine **2** in 81% over two steps (Scheme 1). Advanced kainoid intermediate **6** was prepared according to our previous work^17^ and was condensed with thiosemicarbazide **4** using EDCI to obtain benzylamine **7** in 72%, which was surprisingly stable. *N*-DMB-protected kainoid **7** had to be refluxed in TFA to remove its benzyl group. Full deprotection of the heteroaryl intermediate **8** was accomplished in two steps to afford the desired 2’-aminooxadiazol-4’-yl kainic acid (AODKA, **1**) in 56%. The *syn-C3,C4* stereochemical configuration of **1** at C4 was determined by ^1^H NMR spectroscopy analysis using NOE experiments (see *Supp. Info*.). The heteroaryl’s *exo*-amine is a synthetic handle that can be further functionalized at a late stage. The novel AODKA (**1**) agonist was then applied to investigate the function of kainic acid receptors in astrocytic cells.

## AODKA activates KARs rapidly

To test if AODKA is an effective agonist for KARs, we repeated the phenotypic experiments examining glioexcitability of U118-MG cells. Gratifyingly, cells exposed to 100 μM AOKDA increased their [Ca^2+^]_i_ at a magnitude similar to glutamate (Fig. 4). It also triggered filopodia extension as fast as glutamate (Fig. 4D). Comparing the extent of calcium flux and processes extension in cells shows that AODKA is as effective as glutamate to activate KARs. The action of AODKA on KARs was confirmed by using the antagonist CNQX. Pre-treating the cells with CNQX inhibited filopodiagenesis when AODKA was added to the cells’ medium (Fig. 4B). The selectivity of AODKA for KARs versus other channels remains to be fully evaluated.

In summary, the results presented above demonstrate that KARs contribute directly to the glutamate signaling cascade in U118-MG astrocyte model cells. They also demonstrate that OADKA is a more potent activator of KARs than current common kainoids. We introduced AODKA as a novel and efficient tool to activate KARs which paves the way to exploring the function of these understudied glutamate receptors in the central nervous system.

## METHODS

### Synthesis of AODKA and precursors

#### General Information

NMR spectra were acquired on a 400 MHz Varian NMR AS400 unit equipped with an ATB-400 probe at 25 °C. Infrared spectra (IR) were obtained using a PerkinElmer FT-IR Spectrum Two spectrometer. High resolution mass spectrometry analyses recorded with a HCTultra PTM Discovery System or with a Waters Micromass LCT Premier TOF mass spectrometer. Melting points of solid samples were measured with a IA9200 melting point apparatus (Electrothermal). Column chromatography was carried out on silica gel (230-400 mesh, Silicycle, Quebec).

#### 1-(Isothiocyanatomethyl)-2,4-dimethoxybenzene (3)

To a solution of benzylamine **2** (1.20 g, 7.18 mmol) and triethylamine (1.09 mL, 7.89 mmol) in dichloromethane (50 mL) was added carbon disulfide (4.34 mL, 71.7 mmol). The mixture was stirred at rt for 1 h, and became turbid. To this suspension were added Boc_2_O (1.88 g, 8.61 mmol) and DMAP (87 mg, 0.72 mmol). The solution turned clear and was stirred at rt for an additional 30 mins. The mixture was poured into water and extracted with diethyl ether (3 × 50 mL). The combined organic layer was washed with 5% citric acid aqueous solution, brine, dried over MgSO_4_ and filtered. The filtrate was concentrated under reduced pressure. The residue was purified by column chromatography using ethyl acetate/hexanes as eluent (5-10% gradient). The isocyanate product **3** was recovered as a colourless oil (1.22 g, 81%). ^1^H NMR (400 MHz, CDCl_3_): δ 7.19 (d, *J* = 8.1 Hz, 1H), 6.52 –- 6.44 (m, 2H), 4.60 (s, 2H), 3.84 (s, 5H), 3.82 (s, 3H) ppm. ^13^C NMR (101 MHz, CDCl_3_): δ 161.3, 157.9, 129.4, 115.1, 104.1, 98.6, 55.5, 55.5, 44.1 ppm.

#### *N*-(2,4-Dimethoxybenzyl)hydrazinecarbothioamide (4)

To a solution of isocyanate **3** (987 mg, 4.72 mmol) in isopropanol (10 mL) was added hydrazine hydrate dropwise (432 μL, 60% 5.19 mmol); the addition rapidly caused a white precipitate to form. The suspension was stirred at rt for 20 min. The solid product was recovered by vaccum filtration. It was further washed with ethanol, followed by diethyl ether to afford the thiosemicarbazide **4** (1.13 g, 99%) as a white powder without further purification. FTIR (thin film): 3342, 3280, 3188, 3152, 3004, 2938, 1557 cm^-1^. ^1^H NMR (400 MHz, DMSO-*d*_6_): δ 8.70 (s, 1H), 7.94 (s, 1H), 7.08 (d, *J* = 8.3 Hz, 1H), 6.55 (d, *J* = 2.4 Hz, 1H), 6.46 (dd, *J* = 8.3, 2.3 Hz, 1H), 4.57 (d, *J* = 5.8 Hz, 2H), 4.49 (s, 2H), 3.80 (s, 3H), 3.74 (s, 3H) ppm. ^13^C NMR (101 MHz, DMSO-*d_6_*): δ 181.3, 159.8, 157.8, 129.0, 119.0, 104.2, 98.3, 55.5, 55.2, 41.6 ppm. HRMS (ESI-TOF) m/z: [M+Na]^+^ calcd for C_10_H_15_N_3_O_2_SNa, 264.0777; found, 264.0786.

#### Methyl (*2S,3S,4R*)-4-(5-((2,4-dimethoxybenzyl)amino)-1,3,4-oxadiazol-2-yl)-3-(2-methoxy-2-oxoethyl)-1-((4-nitrophenyl)sulfonyl)pyrrolidine-2-carboxylate (7)

To a solution of acid **3** (369 mg, 0.857 mmol) in dichloromethane (5 mL) was added thiosemicarbazide **4** (310 mg, 1.29 mmol) and EDCI (575 mg, 3.00 mmol). The mixture was stirred at rt for 24 h. The solvent was removed under reduced pressure. The product was purified by column chromatography using a 10-50% gradient of ethyl acetate / hexane as eluent. The oxadiazole product **7** was recovered as a white solid (385 mg, 72%). FTIR (thin film): 3013, 2951, 1742 cm^-1^. ^1^H NMR (400 MHz, CDCl_3_): δ 8.26 (d, *J* = 8.8 Hz, 2H), 7.91 (d, *J* = 8.7 Hz, 2H), 7.20 (d, *J* = 8.2 Hz, 1H), 6.49 (d, *J* = 2.3 Hz, 1H), 6.45 (dd, *J* = 8.2, 2.4 Hz, 1H), 5.27 (s, 1H), 4.42 – 4.21 (m, 3H), 3.91-3.83 (m, 1H), 3.86 (s, 3H), 3.80 (s, 6H), 3.79 – 3.70 (m, 2H), 3.60 (s, 3H), 3.07 (p, *J* = 6.9 Hz, 1H), 2.62 (dd, *J* = 17.4, 8.1 Hz, 1H), 2.35 (dd, *J* = 17.5, 6.8 Hz, 1H) ppm. ^13^C NMR (101 MHz, CDCl_3_): δ 171.3, 171.0, 163.5, 161.1, 158.6, 156.5, 150.3, 143.3, 130.7, 128.4, 124.3, 117.9, 103.8, 98.8, 64.5, 55.5, 55.4, 53.0, 52.0, 50.7, 43.5, 42.6, 37.8, 32.0 ppm. HRMS (ESI-TOF) m/z: [M + Na]^+^ calcd for C_26_H_29_N_5_O_11_SNa, 642.1476; found, 642.1483.

#### Methyl (*2S,3S,4R*)-4-(5-amino-1,3,4-oxadiazol-2-yl)-3-(2-methoxy-2-oxoethyl)-1-((4-nitrophenyl)sulfonyl)pyr-rolidine-2-carboxylate (8)

To the solution of oxadiazole **7** (342 mg, 0.426 mmol) in dichloromethane (5 mL) was added trifluoroacetic acid (0.163 mL, 2.13 mmol). The mixture was stirred at 50 °C. After 3 h, TLC showed full conversion. The solvent was removed under reduced pressure. The product was purified by column chromatography using a 10-50% gradient of ethyl acetate / hexane as eluent. Oxadiazole **8** was recovered as a white solid (181 mg, 90%). FTIR (thin film): 3423, 3363, 3102, 2953, 1733, 1656 cm^-1^. ^1^H NMR (400 MHz, CDCl_3_): δ 8.36 (d, *J* = 8.8 Hz, 2H), 8.01 (d, *J* = 8.8 Hz, 2H), 5.27 (s, 2H), 4.34 (d, *J* = 6.9 Hz, 1H), 3.94 – 3.86 (m, 1H), 3.84 – 3.78 (m, 1H), 3.81 (s, 3H), 3.64 (s, 3H), 3.35 (s, 1H), 3.09 (p, *J* = 6.7 Hz, 1H), 2.59 (dd, *J* = 17.4, 8.3 Hz, 1H), 2.40 (dd, *J* = 17.4, 6.7 Hz, 1H) ppm. ^13^C NMR (101 MHz, CDCl_3_): δ 171.2, 171.0, 150.4, 143.5, 128.6, 124.3, 64.5, 56.0, 53.1, 52.1, 50.5, 42.7, 37.8, 32.1, 29.7 ppm. HRMS (ESI-TOF) m/z: [M+Na]^+^ calcd for C_17_H_19_N_5_O_9_Na, 492.0796; found, 492.0792.

#### Methyl (*2S,3S,4R*)-4-(5-amino-1,3,4-oxadiazol-2-yl)-3-(2-methoxy-2-oxoethyl)pyrrolidine-2-carboxylate (9)

To a solution of oxadiazole **8** (165 mg, 0.351 mmol) in acetonitrile (3 mL) were added thiophenol (54 μL) and potassium carbonate (73 mg). The mixture was stirred at 60 °C for 3 h. The reaction mixture poured into water and extracted with ethyl acetate (3 × 10 mL). The combined organic layer was washed with brine, dried over MgSO_4_ and filtered. The filtrate was concentrated under reduced pressure and the resulting residue was purified by column chromatography using a 5-20% gradient of methanol / dichloromethane as eluent. Oxadiazole amine **9** was recovered as a white solid (67 mg, 67%). FTIR (thin film): 3401, 2931, 2856, 1738, 1659 cm^-1^.^1^H NMR (400 MHz, CDCl_3_): δ 5.51 (s, 2H), 3.76 (s, 3H), 3.72 (d, *J* = 7.9 Hz, 1H), 3.71 – 3.66 (m, 1H), 3.65 (s, 3H), 3.43 (dd, *J* = 11.3, 6.5 Hz, 1H), 3.31 (dd, *J* = 11.3, 4.1 Hz, 1H), 2.97 (dtd, *J* = 9.0, 7.7, 6.0 Hz, 1H), 2.73 (s, 1H), 2.64 (dd, *J* = 17.0, 6.0 Hz, 1H), 2.49 (dd, *J* = 17.0, 9.0 Hz, 1H) ppm. ^13^C NMR (101 MHz, CDCl_3_): δ 174.1, 172.1, 163.2, 159.9, 64.0, 52.5, 51.9, 50.1, 43.5, 39.8, 34.0 ppm. HRMS (ESI-TOF) m/z: [M + H]^+^ calcd for C_11_H_17_N_4_O_5_, 285.1193; found, 285.1194.

#### (*2S,3S,4R*)-4-(5-amino-1,3,4-oxadiazol-2-yl)-3-(carboxymethyl)pyrrolidine-2-carboxylic acid (1)

To a solution of oxadiazole amine **9** (63 mg, 0.22 mmol) in methanol (1 mL) was added LiOH aqueous solution (2.5 M, 2.7 mL). The resulting mixture was stirred at rt for 5 h. The solution was neutralized by addition of 0.5 M hydrochloric acid at 0 °C. The mixture was concentrated under reduced pressure. The residue was dissolved in water (2 mL) and purified by ion-exchange chromatography: ion-exchange resin Dowex 50WX4 100-200 mesh, eluting with 0.5 N aqueous ammonia. The eluting fractions were collected and analyzed by TLC for the presence of the desired product (TLC plates were dried gently with a heat gun before being stained with ninhydrin; further heating revealed the presence of the amino acid **1** as yellow spots.) The fractions containing the product were combined, flash frozen, and the solvents were removed by lyophilization to yield a pale-yellow solid. This product was recrystallized with aqueous ethanol to afford the final product **1** as a white solid (47 mg, 83%). FTIR (thin film): 3395, 2951, 1742, 1727 cm^-1^. ^1^H NMR (400 MHz, D_2_O): δ 4.03 (q, *J* = 6.9 Hz, 1H), 3.93 (d, *J* = 8.9 Hz, 1H), 3.88 (dd, *J* = 12.4, 7.7 Hz, 1H), 3.77 (dd, *J* = 12.5, 5.2 Hz, 1H), 3.08 (td, *J* = 11.3, 10.7, 5.3 Hz, 1H), 2.69 (dd, *J* = 16.8, 4.6 Hz, 1H), 2.09 (dd, *J* = 16.8, 10.8 Hz, 1H) ppm. ^13^C NMR (101 MHz, D_2_O): δ 178.0, 172.4, 164.7, 158.2, 64.4, 46.5, 43.1, 37.7, 36.3 ppm. HRMS (ESI-TOF) m/z: [M+H]^+^ calcd for C_11_H_17_N_4_O_5_, 257.0868; found, 257.0868.

#### Cell culture

Human astrocytoma cell line U118-MG (ATCC, #HTB-15) was gifted by Andis Klegeris, UBC Okanagan. Cells were cultured in Dulbecco’s modified essential medium (DMEM, Gibco 11995-065) supplemented with 10% v/v heat-inactivated fetal bovine serum (HyClone 12483-020) 1% v/v penicillin 10,000 Unit/ml streptomycin 10,000 μg/ml (Gibco 15140-163) and incubated at 37 °C in a humidified atmosphere containing 5% CO_2_.

For experiments, cells at 80-90% confluence were suspended using a pre-warmed 0.25% trypsin-EDTA solution for 5 minutes maximum (Gibco SH30236.02) and transferred to 35 mm glass bottom culture dishes (Mutsunami D1130H) in phenol red-free, high glucose DMEM supplemented with 25 mM HEPES (Gibco 21063-029). Enough cells were transferred to make a ~50% confluent culture-dish after adhesion. The cells were incubated for less than their doubling time (~33 h)^23^ for statistical analysis (see statistics section). This incubation period is long enough for the cells to adopt a flattened shape after adherence which is required to have defined filopodia for measurements.

#### Treatments

On the microscope stage, U118-MG cells were stimulated with glutamate (Alfa Aesar A15031-30), AMPA (Sigma-Aldrich A6816-1MG), kainic acid (Tocris 0222), Ph-KA (made in-house), or AODKA (**1**). The compounds’ stock solutions were prepared freshly in deionized millipore water and filter-sterilized through a 0.22 μm syringe filter. Final concentration of all compounds was 100 μM.

#### Transfection

U118-MG astrocytoma cells were transfected with the F-actin marker mCherry-LifeAct-7 plasmid (gift from Michael Davidson, Addgene #54491) using a calcium phosphate precipitation protocol.^24^ LifeAct-7 is a 17 amino acid peptide that binds to the actin cytoskeleton; its conjugation to mCherry allows one to observe the microfilament rearrangement in active cells. It requires the cellular transcription machinery to be expressed; expression levels may therefore vary between cells.^25^

#### Confocal microscopy and live imaging

The extension of cells’ processes was recorded on an Olympus FV1000 fluorescence confocal microscopy equipped with a Plan-ApoN 60x/i.4 oil-immersion objective. Changes in cell morphology were recorded with 60X objective lenses. To track the processes extension in time, an image was acquired (one second) at 20 seconds intervals. Using intervals and short exposure to laser reduced the photobleaching of the fluorescent signals and maximized cell health during the course of the experiments.

#### Quantification of filopodia extension

The length of twenty filopodia per cell was measured manually using FIJI software. Only filopodia whose entire arbor was captured throughout the imaging period were selected.^26^ A filopodium’s length was measured from the edge of the cell membrane to its apical end. The length of each filopodia was manually measured three times.^27^

To calculate the change in length (Δ length), a process’s length from the image prior to stimulation was subtracted from the length of the same filopodium after 20 minutes. The reported Δ length for each cell is the averaged change in length of the 20 selected filopodia per cell.

#### Time-lapse monitoring: Effect of KA versus AMPA

U118-MG cells transfected with mCherry-LifeAct7 were imaged with an Olympus FV1000 fluorescence confocal microscope equipped with a Plan-ApoN 60X/i.4 oil-immersion objective. Changes in processes extension were monitored in time with images (1 second acquisition) being recorded at 20 seconds intervals. U118-MG cells were monitored for 15 minutes prior to stimulation. The cells were stimulated with AMPA (Sigma-Aldrich A6816-1MG) at a final concentration of 100 μM and monitored for 15 minutes; they were then further stimulated with KA at a final concentration of 100 μM and imaging was pursued for an additional 15 minutes.

#### Statistical analysis

Each value represents the average changes measured from 5 cells, each from a separate culture dish (n = 5); the results are presented as the mean ± S.E.M. Significant differences among groups were determined by two-way analysis of variance (ANOVA) and Sidak’s multiple comparison *post* hoc analysis using GraphPad prism 7.04. A *p* value < 0.05 was considered significant; statistically non-significant data are not starred on the plots.

When testing the compounds, we employed a “completely randomized block design” to minimize technical variations within our samples for ANOVA analysis.^28^ For “completely randomized block design”, cells of a 80-95% confluent dish were trypsinized and re-plated in several 35 mm imaging dishes at a time and incubated in a humidified atmosphere containing 5% CO_2_ at 37 °C. The incubation time is shorter than the doubling time of the U118-MG cells, which allows us to minimize variation of cells within passages before exposing to different treatments.^23,29^

#### Live-cell Ca^2+^ imaging with Fluo4-AM

U118-MG cells grown on a glass-bottom dish were incubated with 1.8 μM Fluo-4 AM (Invitrogen; dissolved in DMSO, then diluted with water to ensure a final concentration of DMSO less than 0.1%) for 10 min in a 5% CO_2_ incubator at 37 °C. Cells were then washed twice with phenol red-free DMEM (Gibco 11995-065) and kept in the same incubator for 20 minutes to allow de-esterification Before imaging, the media was replaced with HBSS upplemented with 2 mM Ca^2+^. Fields of view showing at least 10 cells were acquired using a 20x objective. Results report calcium flux in 106 cells from 6 dishes. Images were recorded at a rate of one frame every 5 seconds using an inverted epifluorescence microscope (Zeiss, Axio ObserverZ.1, controlled by ZEN2 software). Baseline fluorescence was recorded for 60 seconds and averaged (*F_0_*), followed by stimulation with kainoids at a final concentration of 100 μM. ImageJ software (NIH ImageJ, FIJI) was used to correct for background fluorescence for each frame: briefly, pixel intensity values of three background regions were averaged and subtracted from the mean pixel intensity of the whole frame. Changes in Fluo-4 fluorescence (*F*) as a function of time were expressed as *F/F_0_*.

## ASSOCIATED CONTENT

### Supporting Information

Experimental procedures, characterization data, ^1^H and ^13^C NMR spectra. The Supporting Information is available free of charge on the ACS Publications website.

## AUTHOR INFORMATION

### Author Contributions

### Notes

The authors declare no competing financial interest.

## ACKNOWLEDGMENTS

The work was carried out with the support by the Natural Science and Engineering Research Council of Canada (NSERC), the John R. Evans Leaders Fund from the Canadian Fund for Innovation (CFI-JELF), and the University of British Columbia. M.T. and Z.T. gratefully acknowledge UBC for University Graduate Fellowships and a Graduate Dean’s Thesis Fellowship.

## Notes

### Competing Interest Statement

The authors have declared no competing interest.

## REFERENCES

(1) Jin, R.; Banke, T. G.; Mayer, M. L.; Traynelis, S. F.; Gouaux, E. Structural Basis for Partial Agonist Action at Ionotropic Glutamate Receptors. Nat. Neurosci. 2003, 6 (8), 803–810. https://doi.org/10.1038/nn1091.

(2) Cornell-Bell, A. H.; Thomas, P. G.; Caffrey, J. M. Ca2+ and Filopodial Responses to Glutamate in Cultured Astrocytes and Neurons. Can. J. Physiol. Pharmacol. 1992, 70 Suppl, S206–18.

(3) Rose, C. R.; Felix, L.; Zeug, A.; Dietrich, D.; Reiner, A.; Henneberger, C. Astroglial Glutamate Signaling and Uptake in the Hippocampus. Front. Mol. Neurosci. 2018, 10 (January), 1–20. https://doi.org/10.3389/fnmol.2017.00451.

(4) Rojas, H.; Colina, C.; Ramos, M.; Benaim, G.; Jaffe, E. H.; Caputo, C.; DiPolo, R. Na+ Entry via Glutamate Transporter Activates the Reverse Na+/Ca2+ Exchange and Triggers - Induced Ca2+ Release in Rat Cerebellar Type-1 Astrocytes. J. Neurochem. 2007, 100 (5), 1188–1202. https://doi.org/10.1111/j.1471-4159.2006.04303.x.

(5) Araque, A.; Parpura, V.; Sanzgiri, R. P.; Haydon, P. G. Tripartite Synapses: Glia, the Unacknowledged Partner. Trends Neurosci. 1999, 22 (5), 208–215. https://doi.org/10.1016/S0166-2236(98)01349-6.

(6) Hirrlinger, J.; Hülsmann, S.; Kirchhoff, F. Astroglial Processes Show Spontaneous Motility at Active Synaptic Terminals in Situ. Eur. J. Neurosci. 2004, 20 (8), 2235–2239. https://doi.org/10.1111/j.1460-9568.2004.03689.x.

(7) Haber, M.; Zhou, L.; Murai, K. K. Cooperative Astrocyte and Dendritic Spine Dynamics at Hippocampal Excitatory Synapses. J. Neurosci. 2006, 26 (35), 8881–8891. https://doi.org/10.1523/JNEUROSCI.1302-06.2006.

(8) Santello, M.; Toni, N.; Volterra, A. Astrocyte Function from Information Processing to Cognition and Cognitive Impairment. Nat. Neurosci. 2019, 22 (2), 154–166. https://doi.org/10.1038/s41593-018-0325-8.

(9) Bazargani, N.; Attwell, D. Astrocyte Calcium Signaling: The Third Wave. Nat. Neurosci. 2016, 19 (2), 182–189. https://doi.org/10.1038/nn.4201.

(10) Verkhratsky, A.; Kirchhoff, F. Glutamate-Mediated Neuronal-Glial Transmission. In Journal of Anatomy; John Wiley & Sons, Ltd (10.1111), 2007; Vol. 210, pp 651–660. https://doi.org/10.1111/j.1469-7580.2007.00734.x.

(11) Villa, K. L.; Nedivi, E. Previews Glutamate Receptors: Not Just for Excitation. https://doi.org/10.1016/j.neuron.2019.11.025.

(12) Willard, S. S.; Koochekpour, S. Glutamate, Glutamate Receptors, and Downstream Signaling Pathways. Int. J. Biol. Sci. 2013, 9 (9), 948–959. https://doi.org/10.7150/ijbs.6426.

(13) Panatier, A.; Robitaille, R. Astrocytic MGluR5 and the Tripartite Synapse. Neuroscience 2016, 323, 29–34. https://doi.org/10.1016/J.NEUROSCIENCE.2015.03.063.

(14) Matute, C. Therapeutic Potential of Kainate Receptors. CNS Neurosci. Ther. 2011, 17 (6), 661–669. https://doi.org/10.1111/j.1755-5949.2010.00204.x.

(15) Cornell-Bell, A. H.; Prem, T. G.; Smith, S. J. The Excitatory Neurotransmitter Glutamate Causes Filopodia Formation in Cultured Hippocampal Astrocytes. Glia 1990, 3 (5), 322–334. https://doi.org/10.1002/glia.440030503.

(16) Iino, M.; Goto, K.; Kakegawa, W.; Okado, H.; Sudo, M.; Ishiuchi, S.; Miwa, A.; Takayasu, Y.; Saito, I.; Tsuzuki, K.; Ozawa1, S. Glia-Synapse Interaction Through Ca2+-Permeable AMPA Receptors in Bergmann Glia. Science (80-.). 2001, 292 (5518), 926–929. https://doi.org/10.1126/science.1058827.

(17) Tian, Z.; Menard, F. Synthesis of Kainoids and C4 Derivatives. J. Org. Chem. 2018, 83 (11), 6162–6170. https://doi.org/10.1021/acs.joc.8b00179.

(18) Alt, A.; Weiss, B.; Ogden, A. M.; Knauss, J. L.; Oler, J.; Ho, K.; Large, T. H.; Bleakman, D. Pharmacological Characterization of Glutamatergic Agonists and Antagonists at Recombinant Human Homomeric and Heteromeric Kainate Receptors in Vitro. Neuropharmacology 2004, 46 (6), 793–806. https://doi.org/10.1016/j.neuropharm.2003.11.026.

(19) tian, Z.; Tabatabaee, M. S.; Edemann, S.; Gibon, J.; Menard, F. Optical Control of Ca2+-Mediated Morphological Response in Glial Cells with Visible Light Using a Photocaged Kainoid. 2020. https://doi.org/10.26434/CHEMRXIV.11888925.V1.

(20) Tian, Z.; Clark, B. L. M.; Menard, F. Kainic Acid-Based Agonists of Glutamate Receptors: SAR Analysis and Guidelines for Analog Design. ACS Chem. Neurosci. 2019, 10 (10), 4190–4198. https://doi.org/10.1021/acschemneuro.9b00349.

(21) Swanson, G. T.; Sakai, R. Ligands for Ionotropic Glutamate Receptors. Prog. Mol. Subcell. Biol. 2009, 46, 123–157. https://doi.org/10.1007/978-3-540-87895-7_5.

(22) Falcón-Moya1, R.; Sihra, T. S.; Rodríguez-Moreno, A. Kainate Receptors: Role in Epilepsy. Front. Mol. Neurosci. 2018, 11. https://doi.org/10.3389/fnmol.2018.00217.

(23) Westermark, B. The Deficient Density-dependent Growth Control of Human Malignant Glioma Cells and Virus-transformed Glia-like Cells in Culture. Int. J. Cancer 1973, 12 (2), 438–451. https://doi.org/10.1002/ijc.2910120215.

(24) Kingston, R. E.; Chen, C. A.; Rose, J. K. Calcium Phosphate Transfection. Curr. Protoc. Mol. Biol. 2003, 63 (1), 9.1.1–9.1.11. https://doi.org/10.1002/0471142727.mb0901s63.

(25) Riedl, J.; Crevenna, A. H.; Kessenbrock, K.; Yu, J. H.; Neukirchen, D.; Bista, M.; Bradke, F.; Jenne, D.; Holak, T. A.; Werb, Z.; Sixt, M.; Wedlich-Soldner, R. Lifeact: A Versatile Marker to Visualize F-Actin. Nat. Methods 2008, 5 (7), 605–607. https://doi.org/10.1038/nmeth.1220.

(26) Tabatabaee, M. S.; Menard, F. L-Type Voltage-Gated Calcium Channel Modulators Inhibit Glutamate-Induced Morphology Changes in U118-MG Astrocytoma Cells. Cell. Mol. Neurobiol. 2020, 1–9. https://doi.org/10.1007/s10571-020-00828-z.

(27) Zatkova, M.; Reichova, A.; Bacova, Z.; Strbak, V.; Kiss, A.; Bakos, J. Neurite Outgrowth Stimulated by Oxytocin Is Modulated by Inhibition of the Calcium Voltage-Gated Channels. Cell. Mol. Neurobiol. 2018, 38 (1), 371–378. https://doi.org/10.1007/s10571-017-0503-3.

(28) Krzywinski, M.; Altman, N. Comparing Samples—Part II. Nat. Methods 2014, 11 (4), 355–356. https://doi.org/10.1038/nmeth.2900.

(29) Krzywinski, M.; Altman, N. Comparing Samples—Part I. Nat. Methods 2014, 11 (3), 215–216. https://doi.org/10.1038/nmeth.2858.

