## Supplementary information for "Aminooxadiazolyl kainic acid reveals that kainic acid receptors contribute to astrocytoma glutamate signaling"

#### *Electronic Supporting information*

### Chemical Synthesis

**General Synthesis Procedures.** All non-aqueous reactions were carried out under a nitrogen or argon atmosphere, in flame-dried single-neck, round bottom flasks fitted with a rubber septum and with magnetic stirring. Air or water sensitive liquids and solutions were transferred via syringe or stainless-steel cannula. Organic solutions were concentrated by rotary evaporation at 25–45 °C at 50-200 torr. Thin layer chromatography (TLC) was performed on glass plates precoated with Silica gel F254, 250 $\mu$m, 60 Å, from EMD Chemicals Inc (EMD 5715-1). TLC plates were visualized under a 254 or 365 nm UV light source, then stained by chemical reagent: typically, iodine vapors were used first, followed by one of the following solutions: acidic ethanolic vanillin or basic aq. potassium permanganate. The TLC plate was then heated briefly to 200 ºC using a heat gun. Column chromatography purifications were performed with 230-400 mesh silica gel from Silicycle, Quebec (SiliaFlash R12030B, P60, 40-63 $\mu$m, 60 Å).

**Materials.** Reagents and starting materials were purchased from: Sigma-Aldrich, Oakwood Chemicals, Alfa Aesar, Acros Organics, TCI America, or Fisher Scientific and were used as received unless otherwise noted. Tetrahydrofuran, dichloromethane, hexanes, toluene, and diethyl ether were purified on a glass contour solvent purification system under an argon atmosphere.[1] $Et_3N$ and pyridine were distilled from $CaH_2$. Solvents used for chromatographic purifications were obtained from Fischer Scientific or VWR and used without further purification.

**Instruments.** $^1$H, $^{13}$C, and $^{19}$F NMR spectra were recorded on a 400 MHz Varian NMR AS400 equipped with an ATB-400 probe at 25 °C. NMR spectra were analyzed with MestReNova 10 from Mestrelab Research. Proton chemical shifts are expressed in ppm ($\delta$ scale) downfield from tetramethylsilane and are referenced to this standard. Carbon chemical shifts are expressed in ppm ($\delta$ scale) downfield from tetramethylsilane and are referenced to carbon resonance of the NMR solvent ($CDCl_3$ 77.00, *d*$_6$-DMSO 39.52,) or MeOH ($\delta$ 49.50) in case of $D_2O$. Fluorine chemical shifts are expressed in ppm ($\delta$ scale) and referenced to $FCCl_3$ (0.0 ppm). Spectral data are listed as follows: chemical shift, integration, multiplicity (s = singlet, d = doublet, t = triplet, q = quartet, m = multiplet, br s = broad singlet), and coupling constant (*J*, Hz). Infrared spectra (IR) of thin films were obtained using a Spectrum Two FT-IR spectrometer (Perkin-Elmer), and the characteristic absorptions are given in wavenumbers. High-resolution mass spectra were

1 Pangborn, A.B., Giardello, M.A., Grubbs, R.H., Rosen, R.K., and Timmers, F.J.. Safe and convenient procedure for solvent purification. *Organometallics* **1996**, *15*, 1518.

obtained in ESI mode using an HCTultra PTM discovery system spectrometer (Mass Spectrometry Facility, UBC Vancouver). Melting points of solid samples were measured with an IA9200 melting point apparatus (Electrothermal) or micro-melting point apparatus Mel-TempII (Laboratory Devices, USA). The final product was purified by HPLC Waters Delta Prep system on a C18 column (Nova-Pak® HR C18, 19×300mm, 6 $\mu$m, Waters) with fluorimetric detector. Detailed HPLC conditions are described in the corresponding procedures.

### Synthesis

Aminooxazolyl kainic acid (AODKA, **1**) was selected as a class representative for: its strong electron-deficient quadrupole characteristics should enhance π–π stacking interactions with Tyr488 within the kainic acid receptor binding site,[2] and its exocyclic amine allowing for attachment of molecular cargo (e.g., fluorophores, biotin, photoswitch, etc.).

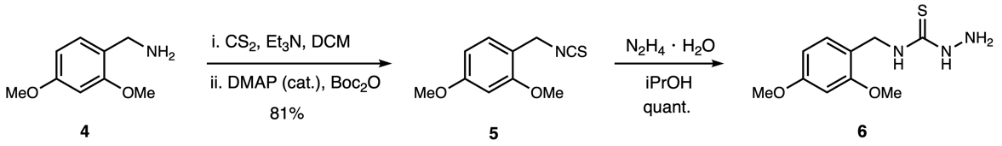


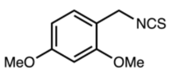


**5**

$C_{10}H_{11}NO_2S$
Molec. weight: 209.26
Exact mass: 209.0510

**1-(Isothiocyanatomethyl)-2,4-dimethoxybenzene (5)**. To a solution of benzylamine **4** (1.20 g, 7.18 mmol) and triethylamine (1.09 mL, 7.89 mmol) in dichloromethane (50 mL) was added carbon disulfide (4.34 mL, 71.7 mmol). The mixture was stirred at rt for 1 h. It resulted in a turbid solution. To this suspension were added $Boc_2O$ (1.88 g, 8.61 mmol) and DMAP (87 mg, 0.72 mmol). The solution turned clear and was stirred at rt for an additional 30 mins. The mixture was poured into water and extracted with ether (3 × 50 mL). The combined organic layer was washed with 5% citric acid aqueous solution, brine, dried over $MgSO_4$ and filtered. The filtrate was concentrated under reduced pressure. The residue was purified by column chromatography using ethyl acetate / hexane as an eluent (5-10% gradient). The isocyanate product **5** was recovered as a colourless oil (1.221 g, 81%). **$^1$H NMR** (400 MHz, $CDCl_3$): δ 7.19 (d, *J* = 8.1 Hz, 1H), 6.52 – 6.44 (m, 2H), 4.60 (s, 2H), 3.84 (s, 5H), 3.82 (s, 3H) ppm. **$^{13}$C NMR** (101 MHz, $CDCl_3$): δ 161.3, 157.9, 129.4, 115.1, 104.1, 98.6, 55.5, 55.5, 44.1 ppm.

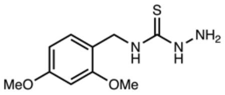


**6**

$C_{10}H_{15}N_3O_2S$
Molec. weight: 241.30
Exact mass: 241.0885

***N*-(2,4-Dimethoxybenzyl)hydrazinecarbothioamide** (**6**). To a solution of isocyanate **5** (987 mg, 4.72 mmol) in isopropanol (10 mL) was added hydrazine hydrate (432 μL, 60%, 5.19 mmol) dropwise, which immediatley formed a white precipitate. The mixture was stirred for 20 min at rt. The solid product was recovered by vaccum filtration. It was further washed by ethanol and ether to afford the thiosemicarbazide **6** (1.132 g, 99%) as a white powder without further purification. **$^1$H NMR** (400 MHz, DMSO-*d*$_6$): δ 8.70 (s, 1H), 7.94 (s, 1H), 7.08 (d, *J* = 8.3 Hz, 1H), 6.55 (d, *J* = 2.4 Hz, 1H), 6.46 (dd, *J* = 8.3, 2.3 Hz, 1H), 4.57 (d,

---

[2] Tian Z, Clark BLM, Menard F. Kainic acid-based agonists of glutamate receptors: SAR analysis and guidelines for analog design. *ACS Chem. Neurosci.* **2019**, 4190.

$J$ = 5.8 Hz, 2H), 4.49 (s, 2H), 3.80 (s, 3H), 3.74 (s, 3H) ppm. **$^{13}$C NMR** (101 MHz, DMSO-$d_6$): δ 181.3, 159.8, 157.8, 129.0, 119.0, 104.2, 98.3, 55.5, 55.2, 41.6 ppm. **FTIR** (thin film): 3342, 3280, 3188, 3152, 3004, 2938, 1557 cm$^{-1}$. **HRMS** (ESI-TOF) m/z: [M + Na]$^+$ calcd for $C_{10}H_{15}N_3O_2SNa$, 264.0777; found, 264.0786.

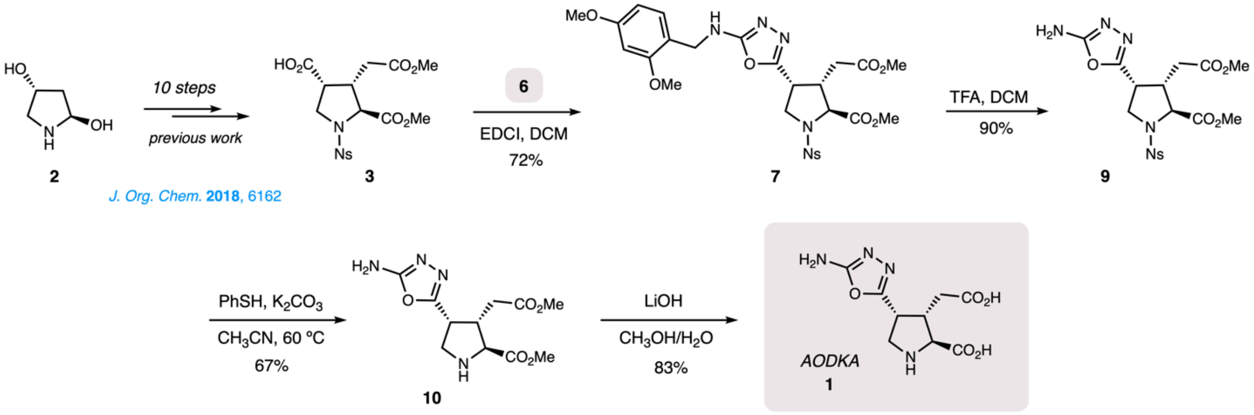


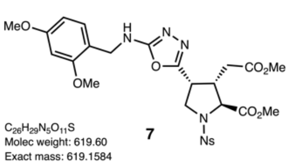


**Methyl (2*S*,3*S*,4*R*)-4-(5'-((*o*,*p*-dimethoxybenzyl)amino)-1',3',4'-oxadiazol-2'-yl)-3-(2''-methoxy-2''-oxoethyl)-1-((*p*-nitrophenyl)sulfonyl)pyrrolidine-2-carboxylate (7).** Advanced kainoid intermediate **3** was prepared from hydroxyprolinol **2** according to the syntheisis reported previously.[3] To a solution of pyrrolidinyl acid **3** (369 mg, 0.857 mmol) in dichloromethane (5 mL) was added thiosemicarbazide **6** (310 mg, 1.29 mmol) and EDCI (575 mg, 3.00 mmol). The mixture was stirred at rt for 24 h. The solvent was removed under reduced pressure. The product was purified by column chromatography using ethyl acetate / hexane as an eluent (10-50% gradient). The oxadiazole product **7** was recovered as a white solid (385 mg, 72%). **FTIR** (thin film): 3013, 2951, 1742 cm$^{-1}$. **$^{1}$H NMR** (400 MHz, $CDCl_3$): δ 8.26 (d, $J$ = 8.8 Hz, 2H), 7.91 (d, $J$ = 8.7 Hz, 2H), 7.20 (d, $J$ = 8.2 Hz, 1H), 6.49 (d, $J$ = 2.3 Hz, 1H), 6.45 (dd, $J$ = 8.2, 2.4 Hz, 1H), 5.27 (s, 1H), 4.42 – 4.21 (m, 3H), 3.91– 3.83 (m, 1H), 3.86 (s, 3H), 3.80 (s, 6H), 3.79 – 3.70 (m, 2H), 3.60 (s, 3H), 3.07 (p, $J$ = 6.9 Hz, 1H), 2.62 (dd, $J$ = 17.4, 8.1 Hz, 1H), 2.35 (dd, $J$ = 17.5, 6.8 Hz, 1H) ppm. **$^{13}$C NMR** (101 MHz, $CDCl_3$): δ 171.3, 171.0, 163.5, 161.1, 158.6, 156.5, 150.3, 143.3, 130.7, 128.4, 124.3, 117.9, 103.8, 98.8, 64.5, 55.5, 55.4, 53.0, 52.0, 50.7, 43.5, 42.6, 37.8, 32.0 ppm. **HRMS** (ESI-TOF) m/z: [M + Na]$^+$ calcd for $C_{26}H_{29}N_5O_{11}SNa$, 642.1476; found, 642.1483.

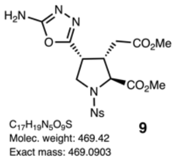


**Methyl (2*S*,3*S*,4*R*)-4-(5'-amino-1',3',4'-oxadiazol-2'-yl)-3-(2''-methoxy-2''-oxoethyl)-1-((*p*-nitrophenyl)sulfonyl)-pyrrolidine-2-carboxylate (9).** To the solution of protected oxadiazole **7** (342 mg, 0.426 mmol) in dichloromethane (5 mL) was added trifluoroacetic acid (0.163 mL, 2.13 mmol). The mixture was stirred at 50 ºC. After 3 h, TLC showed full conversion. The solvent was removed under reduced pressure. The product was purified by column chromatography using ethyl acetate / hexane as an eluent (10-50% gradient). Oxadiazole **9** was recovered as a white solid (181 mg, 90%). **FTIR** (thin film): 3423, 3363, 3102,

[3] Tian Z, Menard F. *Synthesis of kainoids and C4 derivatives*. *J. Org. Chem.* **2018**, *83*, 6162.

2953, 1733, 1656 cm$^{-1}$. **$^{1}$H NMR** (400 MHz, $CDCl_3$): δ 8.36 (d, *J* = 8.8 Hz, 2H), 8.01 (d, *J* = 8.8 Hz, 2H), 5.27 (s, 2H), 4.34 (d, *J* = 6.9 Hz, 1H), 3.94 – 3.86 (m, 1H), 3.84 – 3.78 (m, 1H), 3.81 (s, 3H), 3.64 (s, 3H), 3.35 (s, 1H), 3.09 (p, *J* = 6.7 Hz, 1H), 2.59 (dd, *J* = 17.4, 8.3 Hz, 1H), 2.40 (dd, *J* = 17.4, 6.7 Hz, 1H) ppm. **$^{13}$C NMR** (101 MHz, $CDCl_3$): δ 171.2, 171.0, 150.4, 143.5, 128.6, 124.3, 64.5, 56.0, 53.1, 52.1, 50.5, 42.7, 37.8, 32.1, 29.7 ppm. **HRMS** (ESI-TOF) m/z: [M + Na]$^+$ calcd for $C_{17}H_{19}N_5O_9Na$, 492.0796; found, 492.0792.

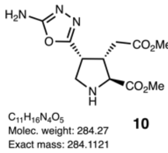


**Methyl (2*S*,3*S*,4*R*)-4-(5'-amino-1',3',4'-oxadiazol-2'-yl)-3-(2''-methoxy-2''-oxoethyl)-pyrrolidine-2-carboxylate** (**10**). To a solution of oxadiazole **9** (165 mg, 0.351 mmol) in acetonitrile (3 mL) were added thiophenol (54 μL) and potassium carbonate (73 mg). The mixture was stirred at 60 °C for 3 h. The reaction mixture poured into water and extracted ethyl acetate (3 × 10 mL). The combined organic layer was washed with brine, dried over $MgSO_4$ and filtered. The filtrate was concentrated under reduced pressure and the resulting residue was by column chromatography using methanol / dichloromethane as an eluent (5-20% gradient). Aminooxadiazole diester **10** was recovered as a white solid (67 mg, 67%). **FTIR** (thin film): 3401, 2931, 2856, 1738, 1659 cm$^{-1}$. **$^{1}$H NMR** (400 MHz, $CDCl_3$): δ 5.51 (s, 2H), 3.76 (s, 3H), 3.72 (d, *J* = 7.9 Hz, 1H), 3.71 – 3.66 (m, 1H), 3.65 (s, 3H), 3.43 (dd, *J* = 11.3, 6.5 Hz, 1H), 3.31 (dd, *J* = 11.3, 4.1 Hz, 1H), 2.97 (dtd, *J* = 9.0, 7.7, 6.0 Hz, 1H), 2.73 (s, 1H), 2.64 (dd, *J* = 17.0, 6.0 Hz, 1H), 2.49 (dd, *J* = 17.0, 9.0 Hz, 1H) ppm. **$^{13}$C NMR** (101 MHz, $CDCl_3$): δ 174.1, 172.1, 163.2, 159.9, 64.0, 52.5, 51.9, 50.1, 43.5, 39.8, 34.0 ppm. **HRMS** (ESI-TOF) m/z: [M + H]$^+$ calcd for $C_{11}H_{17}N_4O_5$, 285.1193; found, 285.1194.

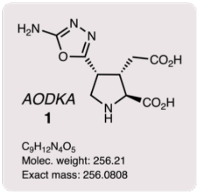


**AODKA: (2*S*,3*S*,4*R*)-4-(5'-Amino-1',3',4'-oxadiazol-2'-yl)-kainic acid** (**1**). To a solution of diester **10** (63 mg, 0.22 mmol) in methanol (1 mL) was added LiOH aqueous solution (2.5 M, 2.7 mL). The resulting mixture was stirred at rt for 5 h. The solution was neutralized by addition of hydrochloric acid (0.5 M) at 0 °C. The mixture was concentrated under reduced pressure. The residue was dissolved in water (2 mL) and purified by ion-exchange chromatography: ion-exchange resin Dowex 50WX4 100-200 mesh, eluting with 0.5 N aqueous ammonia. The eluting fractions were collected and analyzed by TLC for the presence of the desired product. (TLC plates were dried gently with a heat gun before being stained with ninhydrin; further heating revealed the presence of diacid **1** as yellow spots.) The fractions containing the product were combined and flash frozen, and the solvents were removed by lyophilization to yield a pale-yellow solid. This product was recrystallized with aqueous ethanol to afford the final AODKA product **1** as a white solid (47 mg, 83%). **FTIR** (thin film): 3395, 2951, 1742, 1727 cm$^{-1}$. **$^{1}$H NMR** (400 MHz, $D_2O$): δ 4.03 (q, *J* = 6.9 Hz, 1H), 3.93 (d, *J* = 8.9 Hz, 1H), 3.88 (dd, *J* = 12.4, 7.7 Hz, 1H), 3.77 (dd, *J* = 12.5, 5.2 Hz, 1H), 3.08 (td, *J* = 11.3, 10.7, 5.3 Hz, 1H), 2.69 (dd, *J* = 16.8, 4.6 Hz, 1H), 2.09 (dd, *J* = 16.8, 10.8 Hz, 1H) ppm. **$^{13}$C NMR** (101 MHz, $D_2O$): δ 178.0, 172.4, 164.7, 158.2, 64.4, 46.5, 43.1, 37.7, 36.3 ppm. **HRMS** (ESI-TOF) m/z: [M + H]$^+$ calcd for $C_{11}H_{17}N_4O_5$, 257.0868; found, 257.0868.

$^{1}H$ NMR (400 MHz, $CDCl_3$): 1-(Isothiocyanatomethyl)-2,4-dimethoxybenzene (**5**):

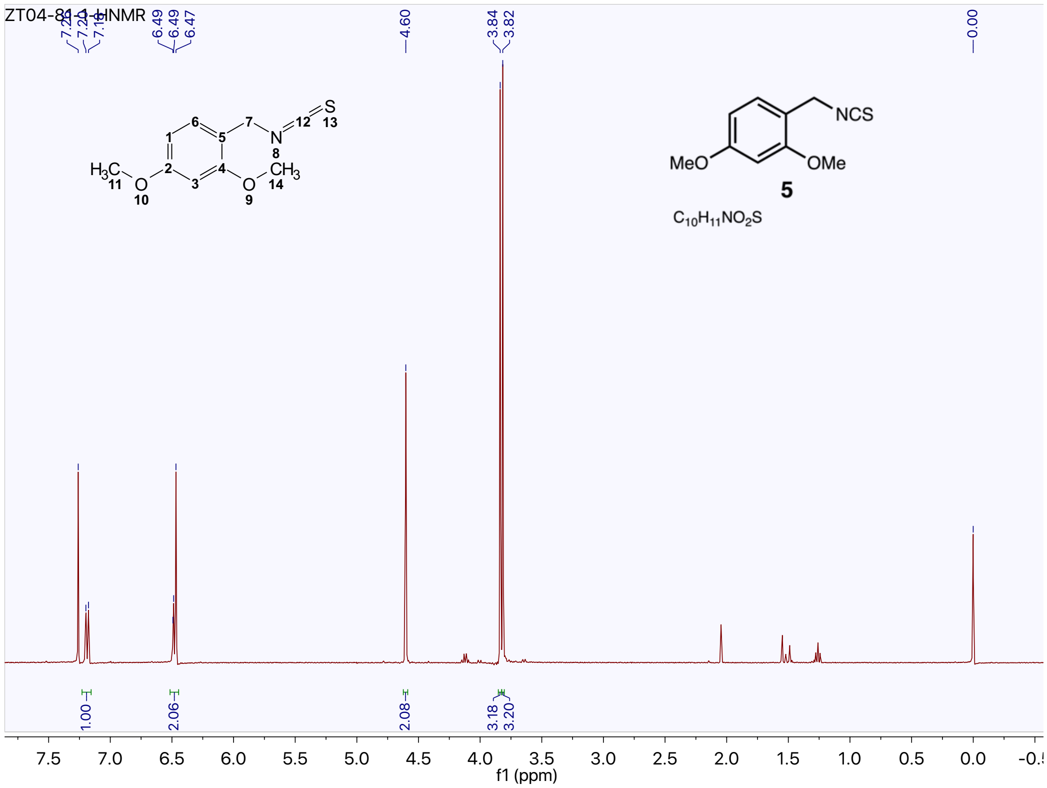


$^{13}C$ NMR (101 MHz, $CDCl_3$): 1-(Isothiocyanatomethyl)-2,4-dimethoxybenzene (**5**).

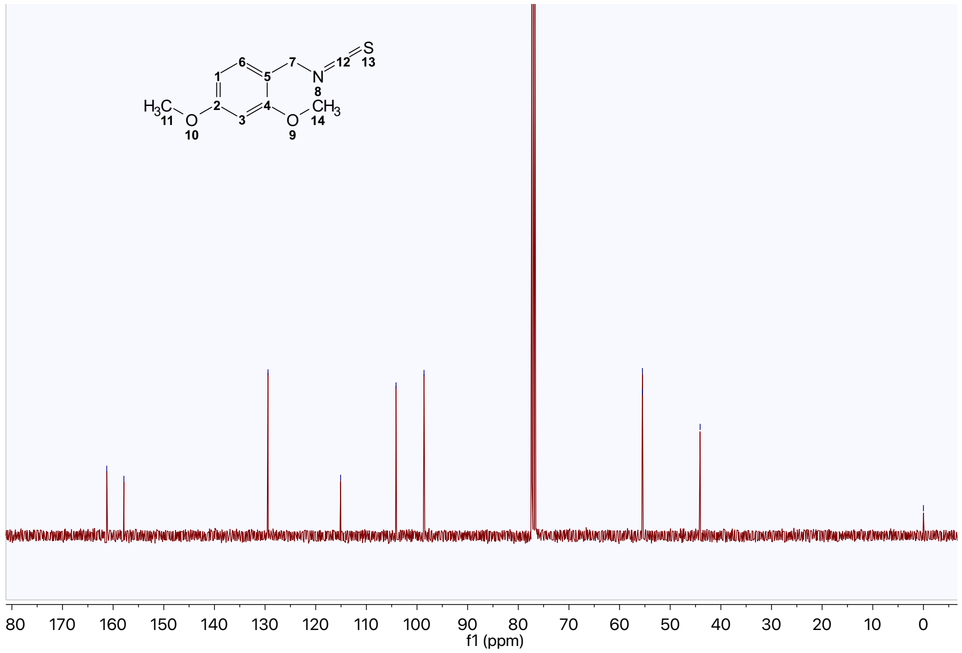

[1]H NMR (400 MHz, DMSO-$d_6$): *N*-(2,4-Dimethoxybenzyl)hydrazinecarbothioamide (**6**):

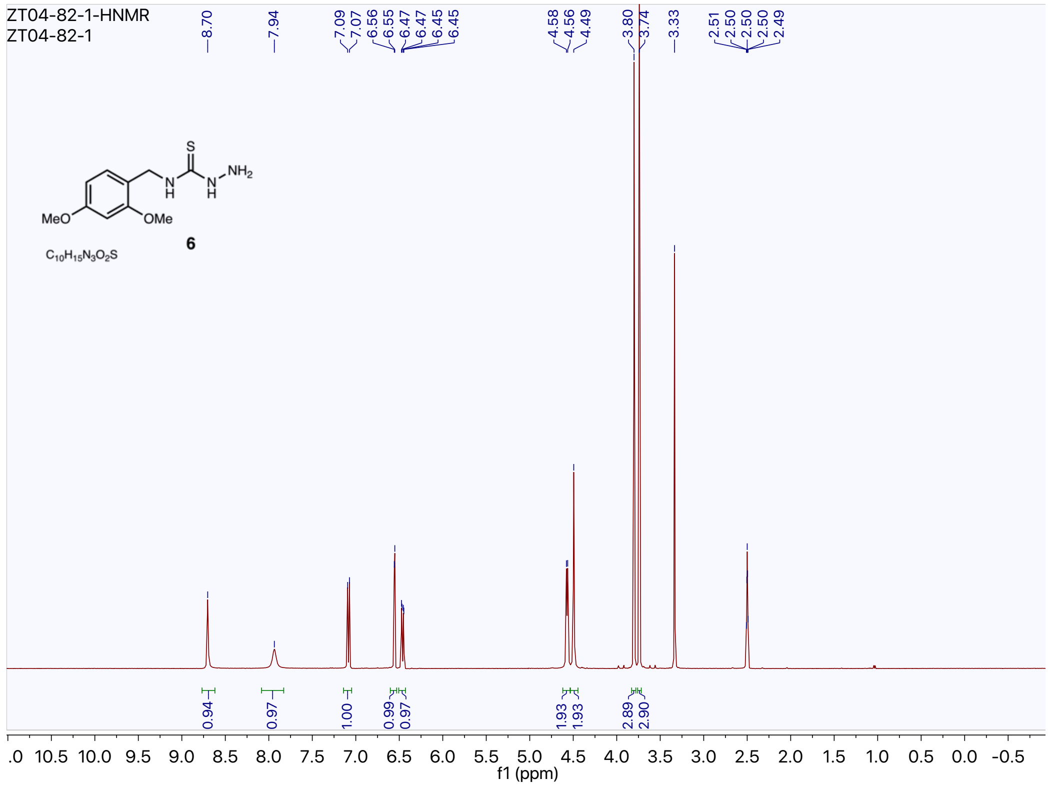


[13]C NMR (101 MHz, DMSO-$d_6$): *N*-(2,4-Dimethoxybenzyl)hydrazinecarbothioamide (**6**):

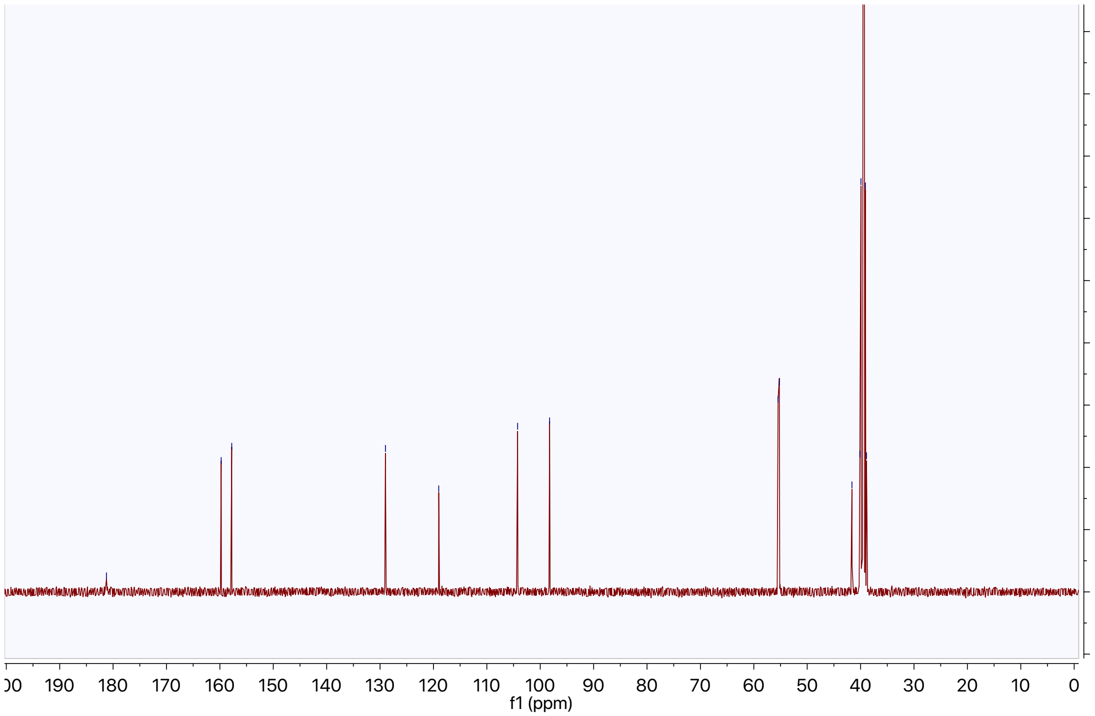

[1]H NMR (400 MHz, $CDCl_3$): Methyl (2*S*,3*S*,4*R*)-4-(5-((2,4-dimethoxybenzyl)amino)-1,3,4-oxadiazol-2-yl)-3-(2-methoxy-2-oxoethyl)-1-((4-nitrophenyl)sulfonyl)pyrrolidine-2-carboxylate (**7**):

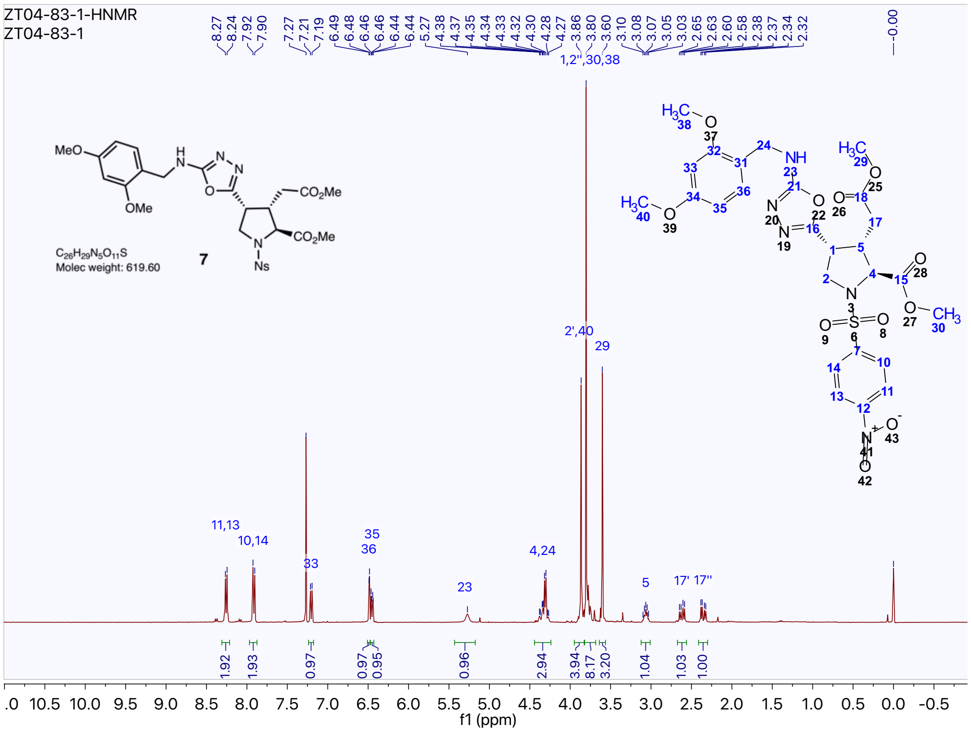

[13]C NMR (101 MHz, $CDCl_3$): Methyl (2*S*,3*S*,4*R*)-4-(5-((2,4-dimethoxybenzyl)amino)-1,3,4-oxadiazol-2-yl)-3-(2-methoxy-2-oxoethyl)-1-((4-nitrophenyl)sulfonyl)pyrrolidine-2-carboxylate (**7**):

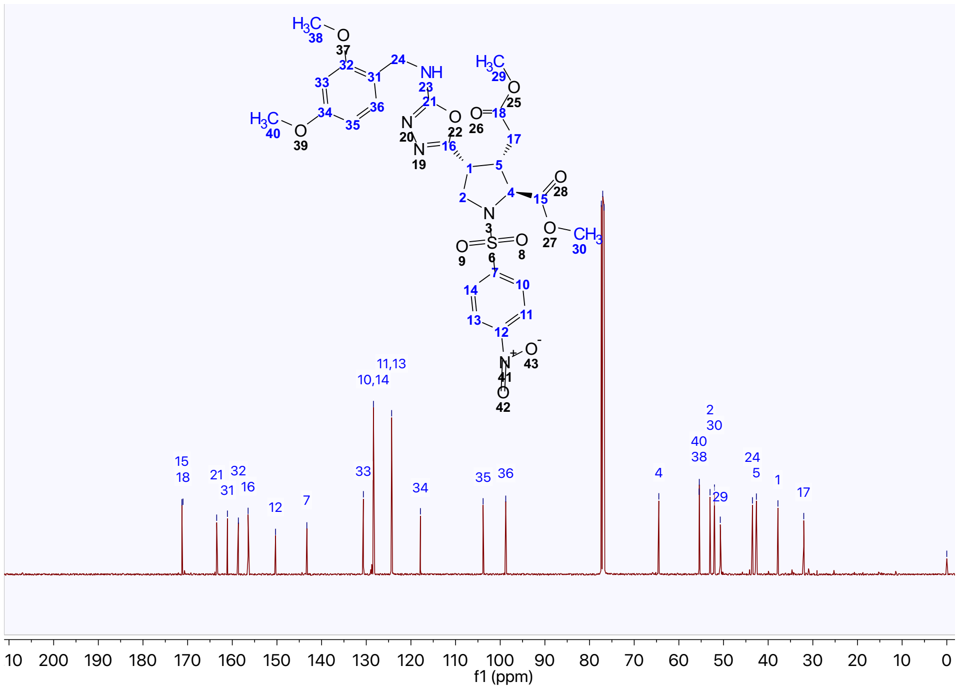

[1]H NMR (400 MHz, $CDCl_3$): Aminooxadiazolyl-(*N*-nosyl)kainic acid diester (**9**):

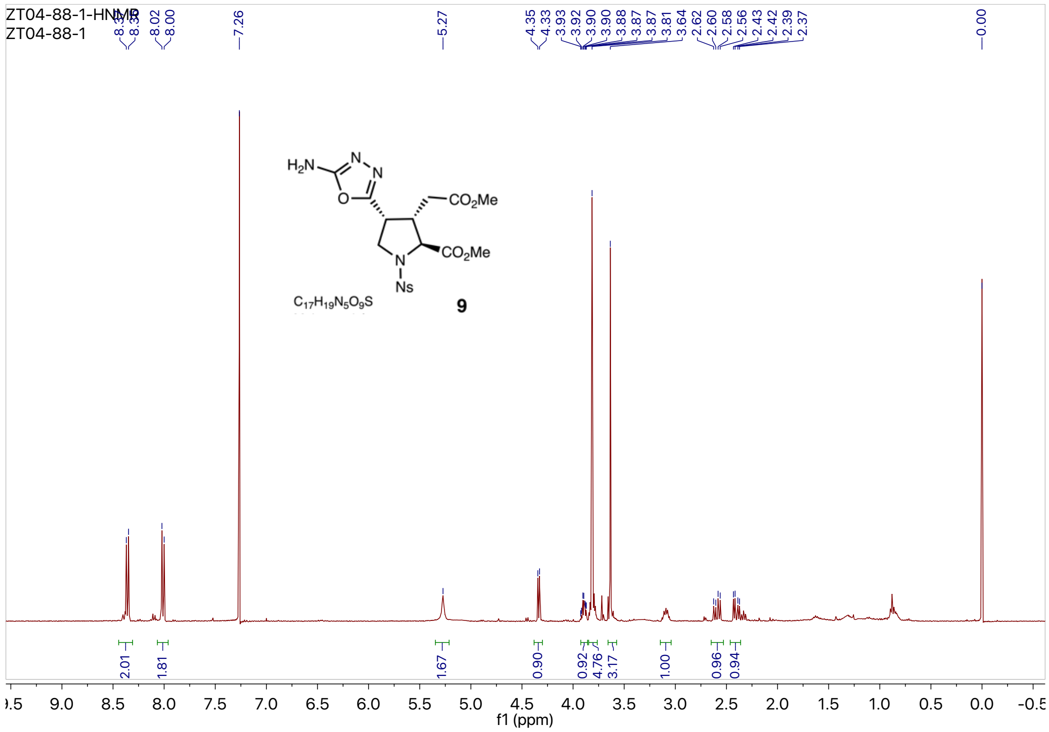


[13]C NMR (101 MHz, $CDCl_3$): Aminooxadiazolyl-(*N*-nosyl)kainic acid diester (**9**):

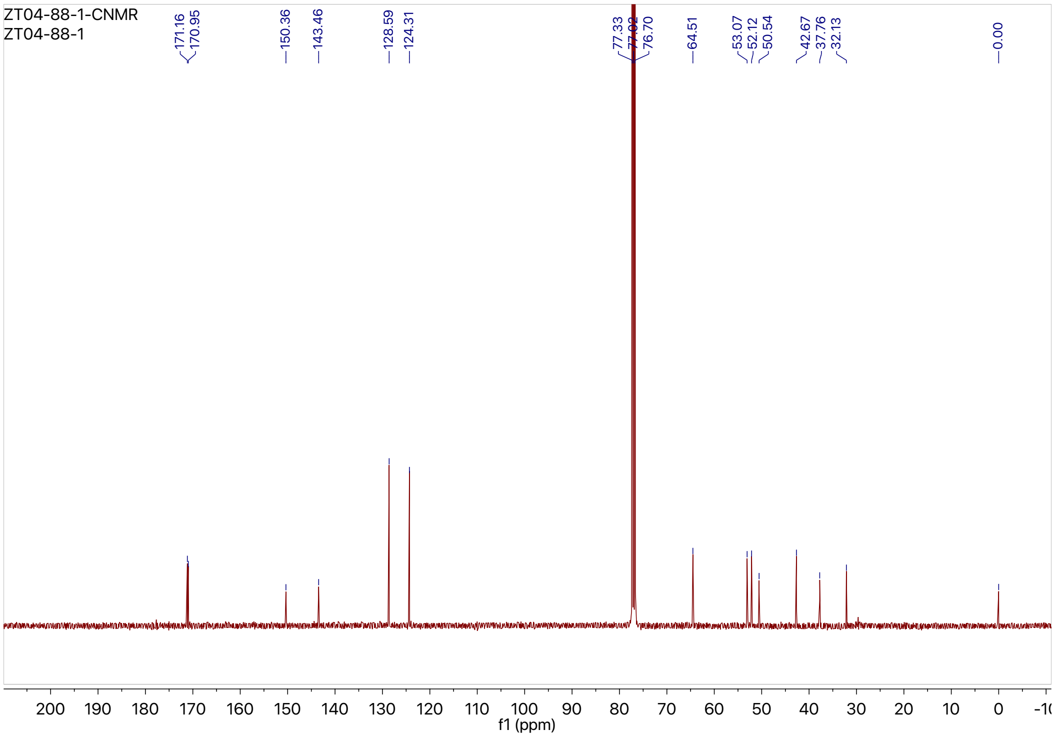

$^{1}H$ NMR (400 MHz, $CDCl_3$): Aminooxadiazolyl-kainic acid diester (**10**):

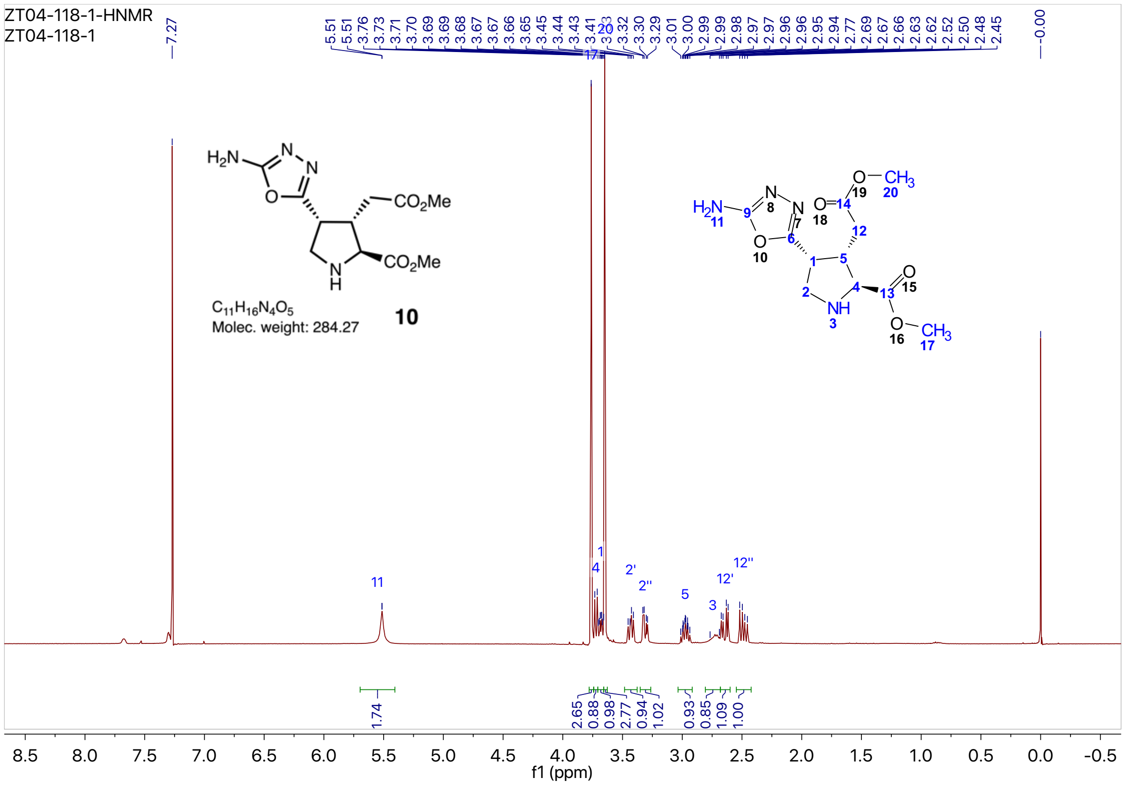


$^{13}C$ NMR (101 MHz, $CDCl_3$): Aminooxadiazolyl-kainic acid diester (**10**):

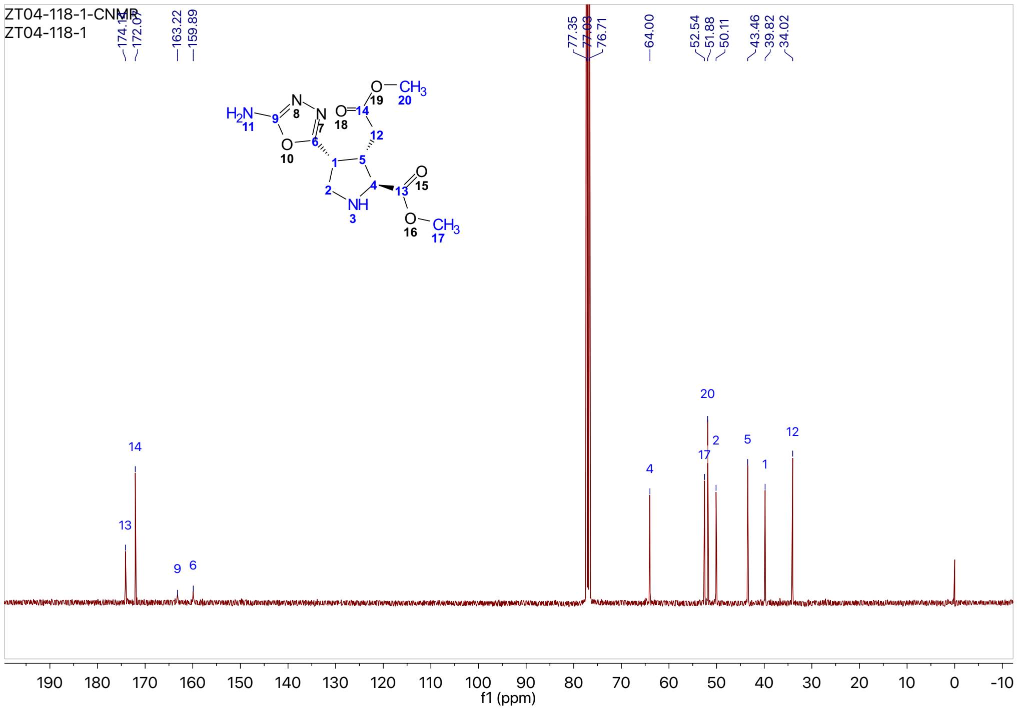

$^{1}H$ NMR (400 MHz, $D_2O$): Aminooxadiazolyl-kainic acid, AODKA (**1**):

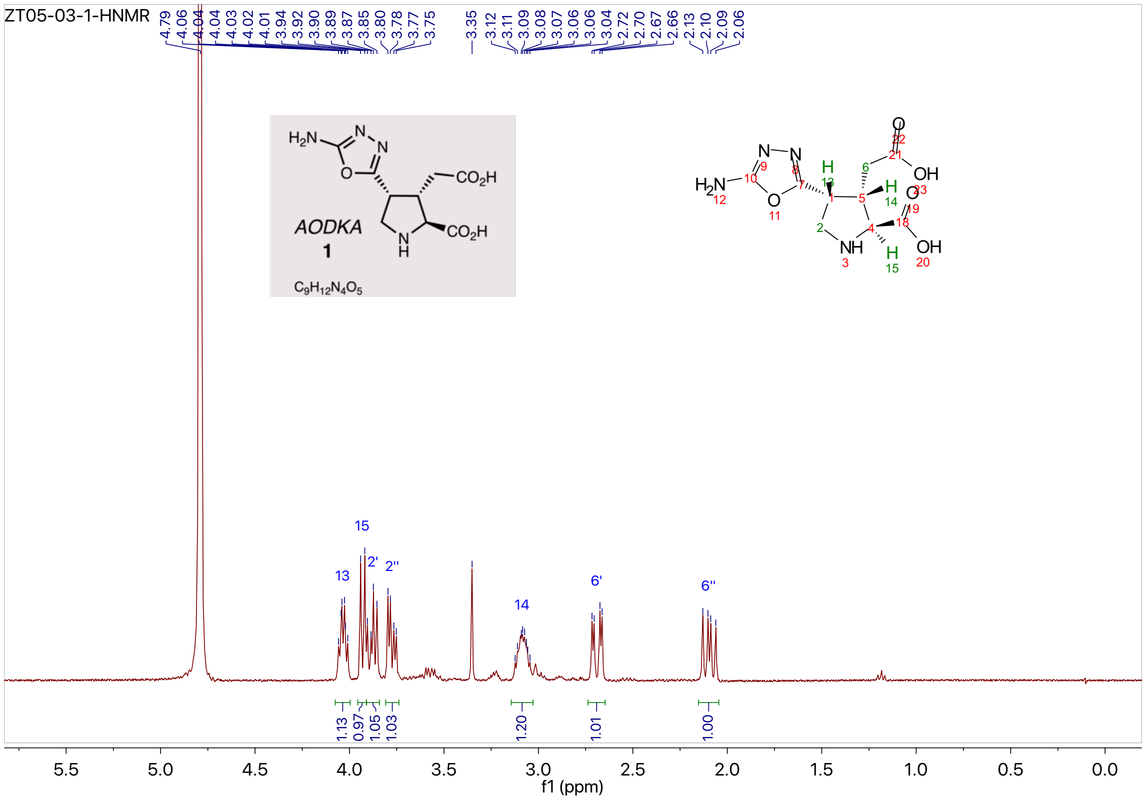

$^{13}C$ NMR (101 MHz, $D_2O$): Aminooxadiazolyl-kainic acid, AODKA (**1**):

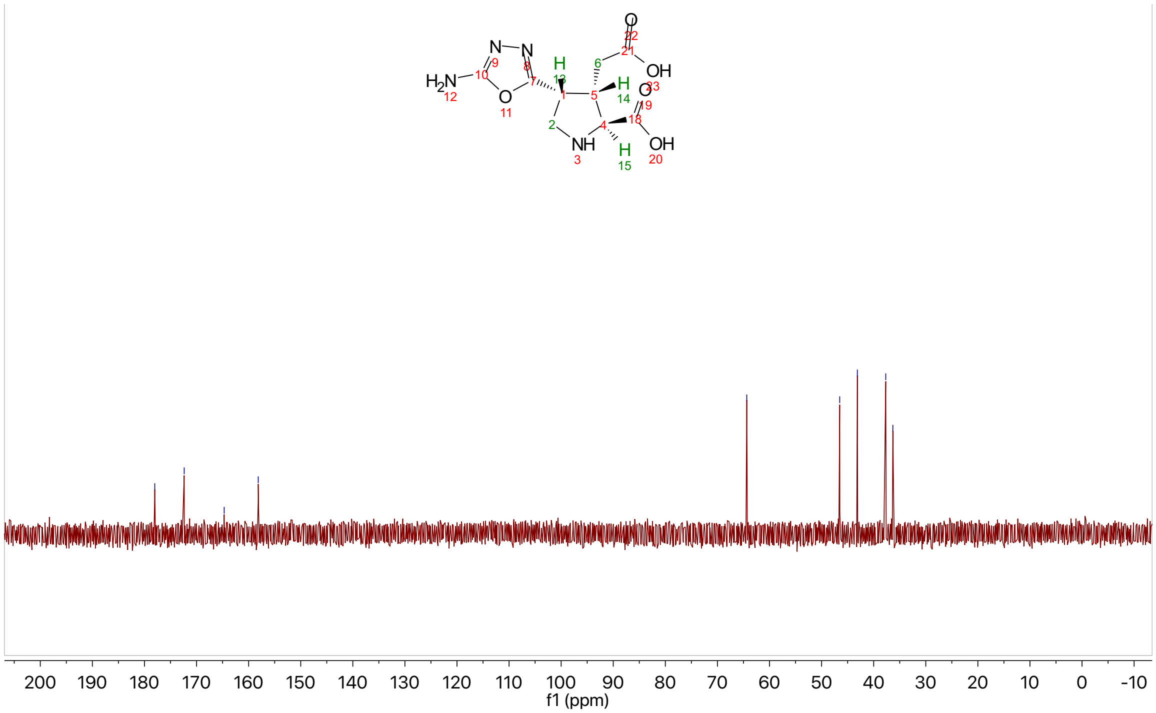
